# A constrained SSU-rRNA phylogeny reveals the unsequenced diversity of photosynthetic Cyanobacteria (Oxyphotobacteria)

**DOI:** 10.1101/315697

**Authors:** Luc Cornet, Annick Wilmotte, Emmanuelle J. Javaux, Denis Baurain

**Affiliations:** InBioS – PhytoSYSTEMS, Eukaryotic Phylogenomics, University of Liège, Belgium; UR Geology – Palaeobiogeology-Palaeobotany-Palaeopalynology, University of Liège, Belgium; InBioS – CIP, Centre for Protein Engineering, University of Liège, Belgium; BCCM/ULC collection of cyanobacteria, University of Liège, Belgium

**Keywords:** Cyanobacteria, Phylogeny, Diversity, Genomics, SSU rRNA (16S), Sequenced genome fraction

## Abstract

**Objectives:** Cyanobacteria are an ancient phylum of prokaryotes that contain the class Oxyphotobacteria, the unique bacterial group able to perform oxygenic photosynthesis. This group has been extensively studied by phylogenomics during the last decade, notably because it is widely accepted that Cyanobacteria were responsible for the spread of photosynthesis to the eukaryotic domain. The aim of this study was to evaluate the fraction of the oxyphotobacterial diversity for which sequenced genomes are available for genomic studies. For this, we built a phylogenomic-constrained SSU rRNA (16S) tree to pinpoint unexploited clusters of Oxyphotobacteria that should be targeted for future genome sequencing, so as to improve our understanding of Oxyphotobacteria evolution.

**Results:** We show that only a little fraction the oxyphotobacterial diversity has been sequenced so far. Indeed 31 rRNA clusters on the 60 composing the photosynthetic Cyanobacteria have a fraction of sequenced genomes <1%. This fraction remains low (min = 1%, median = 11.1 %, IQR = 7.3) within the remaining “‘sequenced” clusters that already contain some representative genomes. The “unsequenced” clusters are scattered across the whole Oxyphotobacteria tree, at the exception of very basal clades (G, F, E) and the Oscillatoriales clade (A), which have higher fractions of representative genomes. Yet, the very basal clades still feature some (sub)clusters without any representative genome. This last result is especially important, as these basal clades are prime candidate for plastid emergence.

## Introduction

Oxyphotobacteria are a class of the phylum Cyanobacteria, one of the most important group of prokaryotes [1]. This morphologically diversified class is the only group of Bacteria that can carry out oxygenic photosynthesis. It is commonly assumed that this bioenergetic process appeared in Oxyphotobacteria (or their ancestor) and caused the Great Oxidation Event (GOE) 2.4 billion years ago [2–5]. The atmospheric increase in free oxygen had a critical impact on early Earth evolution, including on Oxyphotobacteria themselves, with the appearance of new morphological features (e.g., akinetes) [6]. Oxyphotobacteria were also responsible for the spread of photosynthesis to eukaryotic lineages, through an initial event of primary endosymbiosis, followed by higher-order endosymbioses [7]. In consequence of their great scientific interest, numerous phylogenies of Oxyphotobacteria have been published during the last decade (eg, [5, 8–19]), using ever more genomes as it appears within public repository. The diversity within Oxyphotobacteria is very high, both morphologically and genetically, and their GC-content ranges from 35 to 71 [20]. Therefore, it is important to assess how well the existing genomes represent the whole diversity of the class and detect the lineages for which representative genomes are not yet available. Indeed, only 659 genomes of Oxyphotobacteria have been sequenced, whereas 6997 non-redundant rRNA sequences can be found in the SILVA database (as of October 2017). In this short communication, we built a SSU rRNA (16S) phylogeny to investigate the distribution of representative genomes across oxyphotobacterial diversity, so as to guide the selection of taxa for future sequencing projects.

## Main text

We took advantage of 7603 different SSU rRNA (16S) sequences [6997 from SILVA 128 Ref NR [21], 466 from genome assemblies publicly available on the NCBI [22] and 140 from the BCCM/ULC public collection of Cyanobacteria (http://bccm.belspo.be/about-us/bccm-ulc)], leading to a dataset of 2302 unique Operational Taxonomic Units (OTUs) after dereplication at an identity threshold of 97.5%. Because multiple sequence alignment (MSA) quality is essential in single-gene analyses [23–25], we used an in-house hand-curated MSA as a starting point to carefully align all the OTUs. This MSA represented a high fraction of known Oxyphotobacteria diversity, as it completely included the broadly sampled dataset of Schirrmeister et al. (2011) [26]. To alleviate the stochastic error plaguing single-gene analyses [27], we constrained our SSU rRNA (16S) phylogeny using a more reliable multi-gene phylogeny computed on a subset of the OTUs at hand. To this end, we used the phylogeny of Oxyphotobacteria developed in Cornet et al. (2018b) [28], the largest ever-built in terms of conserved sites (>170,000 unambiguously aligned amino-acid positions from 675 genes). This strategy allowed us to easily compare the genome diversity available for phylogenomic studies to the more completely sampled rRNA diversity of Oxyphotobacteria.

As shown in Figure 1, 31 rRNA clusters have a fraction of sequenced genomes <1% and are thus nearly completely ignored in phylogenomic studies. These “unsequenced” clusters are scattered across the whole oxyphotobacterial tree, at the exception of very basal clades (G, F, E) and the Oscillatoriales clade (A), which have higher fractions of representative genomes. The group C (C1, C2, C3) appears paraphyletic in our tree, and is especially rich in “unsequenced” clusters (14 out of 31), including the largest one (212 OTUs). In all the remaining “‘sequenced” clusters that contain representative genomes, the fraction of sequenced genomes remains low (min = 1%, median = 11.1 %, IQR = 7.3). It is thus clear that genomes have not been evenly sampled across the tree. To help researchers to select organisms for future sequencing projects, our final dataset and tree are made available (see below. In these files, SSU rRNA (16S) genes can be readily identified by their accession number, whether for SILVA/ULC strains or for rRNA genes predicted from genome assemblies.

**Figure 1:**
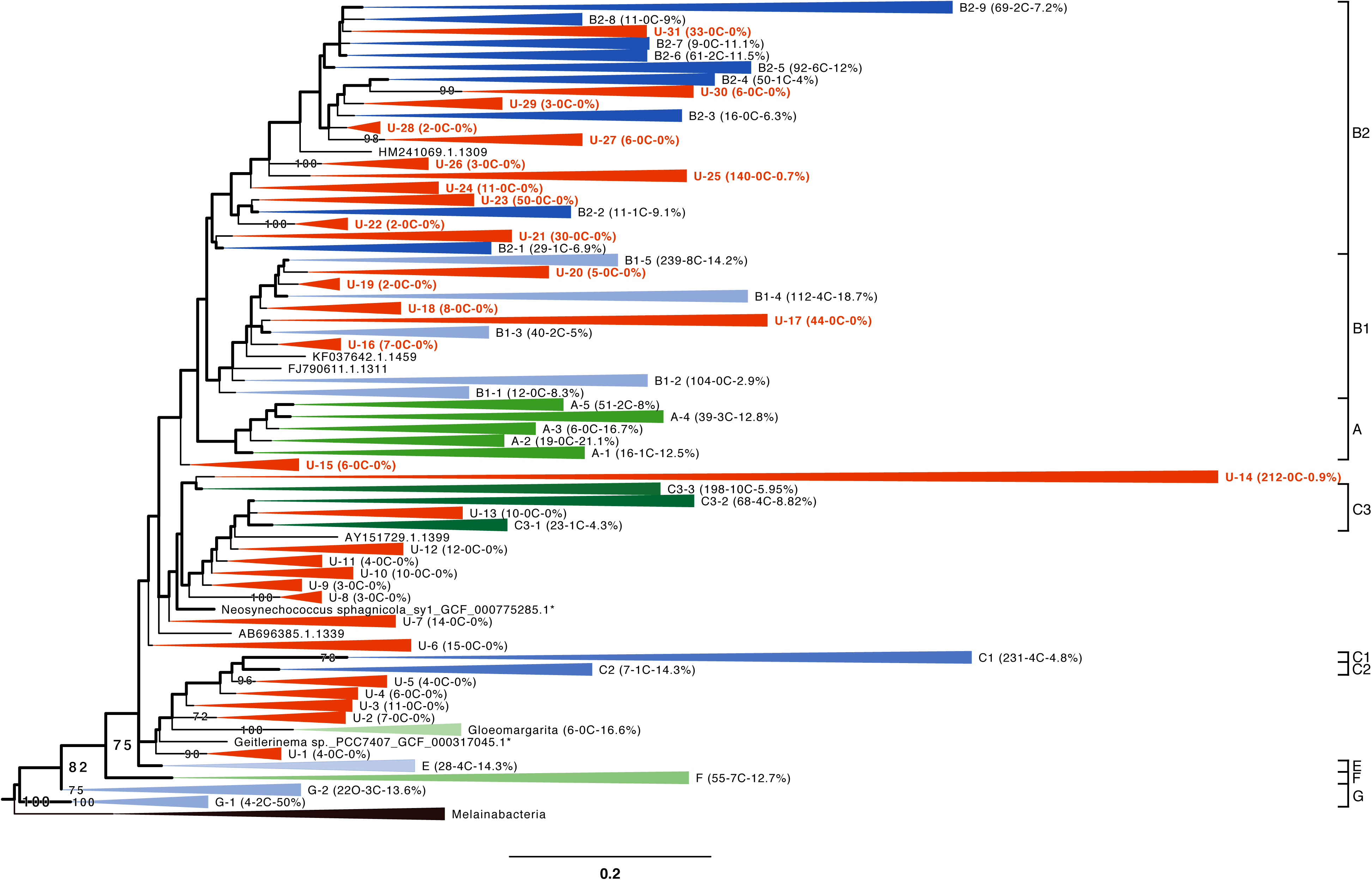
Collapsed constrained phylogeny of Oxyphotobacteria based on an alignment of 2302 SSU rRNA (16S) sequences (1233 unambiguously aligned nucleotide positions). The backbone of the tree shown in thick lines corresponds to the branches constrained by a phylogenomic tree of 71 OTUs x 170,983 positions (Cornet et al., 2018b). Clusters on a red background have a fraction of sequenced genomes <1% and can be considered as “unsequenced” due to the lack of representative genomes. Clusters on another colored background have a higher fraction of sequenced genomes (“sequenced” clusters) and were annotated using the clade names of Shih et al. (2013). For each cluster (at the exception of Melainabacteria), we report in brackets the number of OTUs, the number of constrained genome sequences and the fraction of sequenced genomes.

The emergence of eukaryotic plastids from Oxyphotobacteria has been the subject of different hypotheses during the last decade, the most frequent being an early origin around clade E, composed of *Acaryochloris* and *Thermosynechococcus* species [17, 18]. By focusing on the base of our oxyphotobacterial phylogeny, Figure 2 shows that in spite of the presence of a number of representative genomes, very basal clades (G, F, E) still feature some clusters without any genome (highlighted in yellow in Figure 2). Therefore, these clusters should be targeted in priority for genome sequencing if one wants to resolve the early diversification of Oxyphotobacteria and the origin of plastids by assembling phylogenomic datasets more faithful to the true diversity.

**Figure 2:**
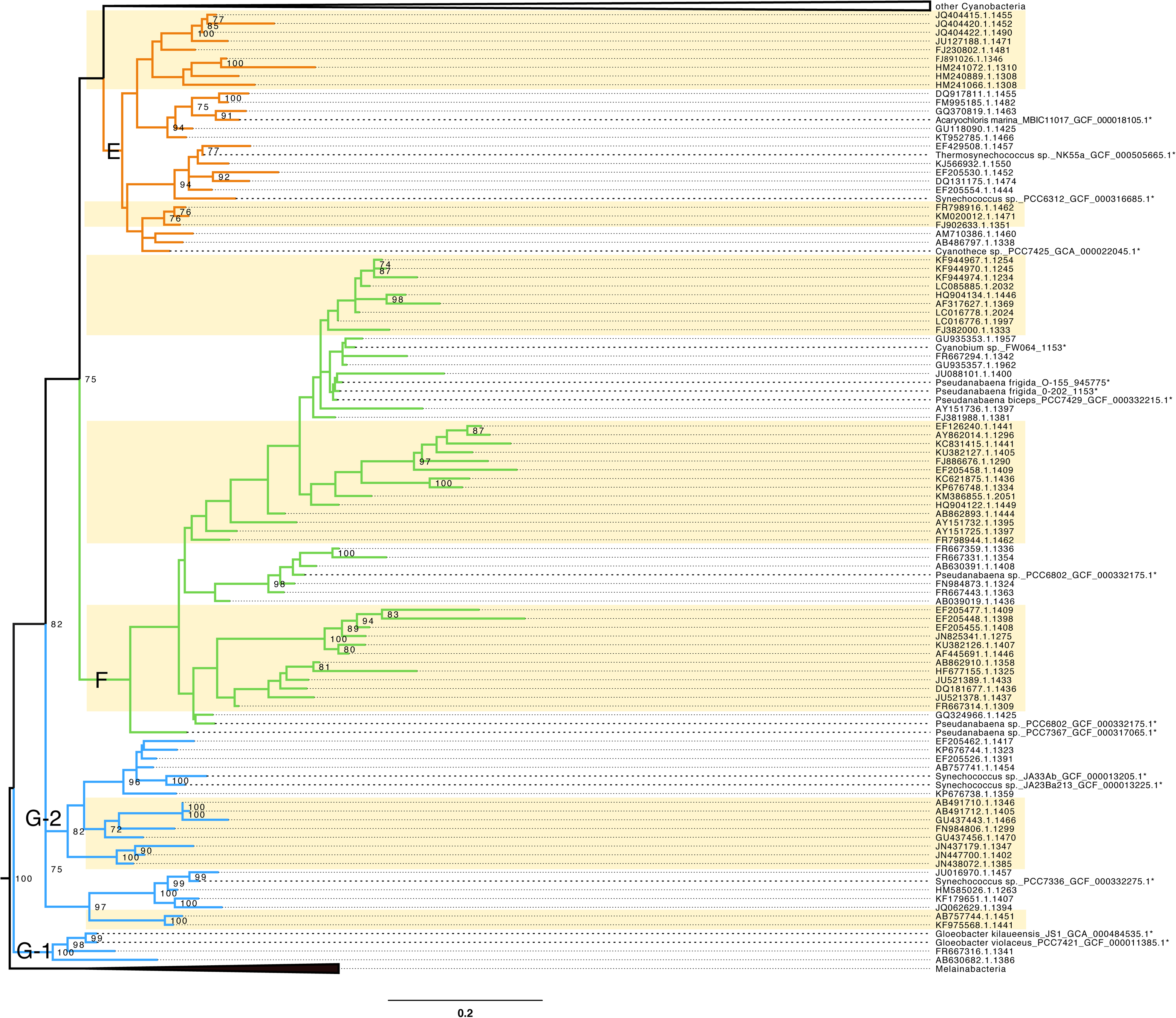
Expanded basal part of the constrained rRNA phylogeny of Oxyphotobacteria, with clusters E, F, G1 and G2. (*) show constrained genome sequences. The color of the backbone corresponds to the groups of Shih et al. (2013). “Unsequenced” clusters (both monophyletic and paraphyletic) are shown on a yellow background.

## Methods

Cyanobacterial (651 Oxyphotobacteria and 8 Melainabacteria) genome assemblies were downloaded from the NCBI FTP server on October 1^st^, 2017. SSU rRNA (16S) genes were predicted for all assemblies with RNAmmer v1.2 [29]. Predicted rRNA genes were taxonomically classified with SINA v1.2.11 [30], using the release 128 of SILVA database composed of 1,922,213 SSU rRNA (16S) reference sequences [21]. Among the 651 Oxyphotobacteria assemblies, 193 were devoid of Oxyphotobacteria rRNA genes (176 without any prediction and 17 with only non-cyanobacterial predicted genes). All Oxyphotobacteria rRNA genes (7001 sequences) of SILVA 128 Ref NR (non-redundant) were downloaded on October 1^st^, 2017. Four SSU rRNA sequences with a SILVA-provided incongruent lineage (e.g., Proteobacteria for *Prochlorothrix hollandica* PCC 9006) were deleted from the dataset, whereas the 466 sequences predicted with RNAmmer from genome assemblies were added, along with 140 new sequences from strains of the BBMC/ULC Cyanobacteria collection (http://bccm.belspo.be/about-us/bccm-ulc), hence leading to a total of 7603 SSU rRNA (16S) sequences (including 8 Melainabacteria).

The dataset was dereplicated with CD-HIT (cd-hit-est) v4.6 [31], using an identity threshold of 97.5%, an alignment coverage control of 0.0 for the longest sequence and 0.9 for the shortest sequence. The value of 97.5% for the identity threshold was chosen by following the OTU definition of Taton et al. (2003) [32]. It probably underestimates the true number of different species, since Edgar (2018) [33] has recently shown that the bacterial species delineation threshold is closer to 99% for full-length sequences. However, for the purpose of this study aimed at pinpointing large patches of unsequenced diversity, a coarse-grained clustering was sufficient. The cluster file produced by CD-HIT was retrieved and 13 representative sequences initially selected by CD-HIT were replaced by genomic sequences from the same CD-HIT clusters, so as to be able to enforce the phylogenomic constraints from Cornet et al. (2018b) [28] when inferring the rRNA tree. This process led to a dataset of 2302 OTUs.

The 2302 sequences were aligned using the BLASTN-based software “two-scalp” (D. Baurain; https://metacpan.org/release/Bio-MUST-Apps-TwoScalp), starting from a hand-curated MSA (1252 sites post-BMGE; 1.52% missing character states) of 1128 SSU rRNA (16S) sequences (L. Cornet; unpublished). The 1128 sequences had been selected in order to be as close as possible to the dataset of Schirrmeister et al. (2011) [26], which represented all the Oxyphotobacteria diversity known at that time. Unambiguously aligned sites were selected with BMGE v1.12 [34], using moderately severe settings (entropy cut-off 0.5, gap cut-off 0.2) and yielding a high-quality MSA of 1233 conserved sites with only 1.83% missing character states. In comparison, building a MSA from scratch on the same dataset with MAFFT v7.273 [35], using its automatic mode, led to only 1013 conserved sites (2.52% missing character states) after applying the same protocol to select the unambiguously aligned sites.

The tree was inferred with RAxML v8.1.17 [36] under a GTR+Γ4 model with a 100x rapid bootstrap analysis, using a constraining tree of 75 different Oxyphotobacteria. The constraining tree was a phylogenomic tree based on a concatenation of 675 genes (>170,000 unambiguously aligned amino-acid positions; [28]). However, four organisms (*Leptolyngbya* sp. GCF_000733415.1, Oscillatoriales cyanobacterium GCF_000309945.1, *Leptolyngbya* sp. OTC1/1, *Phormidium priestleyi* ANT.PROGRESS2.5) had to be pruned before inference due to the lack of genuine rRNA genes in their assemblies. The rRNA tree was then manually collapsed using FigTree v1.4.3 (http://tree.bio.ed.ac.uk/software/figtree/) and further arranged in InkScape v0.92 [37]. For each collapsed subtree, a “sequenced genome fraction” statistic was computed by taking the ratio between the number of OTUs having a publicly available genome in their CD-HIT cluster to the total number of OTUs in the subtree.

## Limitations

Among the 659 available Oxyphotobacteria genomes, 193 genomes were either completely devoid of rRNA genes (or only featured taxonomically incongruent rRNA genes) and were thus unusable for phylogeny (see Methods for details). Obviously, the results of this study might have been different if we had used these genomes to evaluate the sequenced genome fraction. The absence of rRNA genes is explained by the fact that the majority of these genomes (143 out of 193) were assembled with a metagenomic pipeline, according to NCBI metadata. This phenomenon was noticed in Cornet et al. (2018a) [38] and is due to the frequent loss of rRNA sequences during metagenomic assembly [28]. Given that neither the isolation source nor the assembly pipeline were easy to determine, we decided not to consider these 193 genomes in our analysis by precautionary principle. Hence, a significant number of these metagenomes were found in the genome section of RefSeq, even though it is not allowed. This decision is moreover consolidated by the observation that only a small number (about 20) of SSU rRNA (16S) sequences could be recovered after extensive searches on the NCBI using the “strain” names of these 193 genomes.

## List of abbreviations

OTUs: Operational Taxonomic Units
MSA: Multiple sequence alignment

## Declarations

### Author contributions

LC and DB designed the experiments, LC performed the computational analyses and drew the figures, LC and DB wrote the manuscript, with the assistance of AW and EJJ. All authors read and approved the final manuscript.

## Acknowledgment

We thank Yannick Lara (ULiège) for the cultivation and sequencing of some the BCCM/ULC strains. More generally, we are grateful to the BCCM/ULC public collection of Cyanobacteria for sharing the SSU rRNA (16S) sequences of 140 strains. Rosa Gago is acknowledged for her help when drawing the figures.

## Availability of data and materials

The dataset (final MSA and ML tree) supporting the conclusions of this article is available as compressed archive in the figshare repository through the following link [https://doi.org/10.6084/m9.figshare.6225641]. All intermediate files are also available upon request.

## Consent to publish

Not applicable

## Ethics approval and consent to participate

Not applicable

